# Genome sequence and comparative analysis of reindeer (*Rangifer tarandus*) in northern Eurasia

**DOI:** 10.1101/739995

**Authors:** Melak Weldenegodguad, Kisun Pokharel, Yao Ming, Mervi Honkatukia, Jaana Peippo, Tiina Reilas, Knut H. Røed, Juha Kantanen

## Abstract

Reindeer are semi-domesticated ruminants that have adapted to the challenging northern Eurasian environment characterized by long winters and marked annual fluctuations in daylight. We explored the genetic makeup behind their unique characteristics by *de novo* sequencing the genome of a male reindeer and conducted gene family analyses with nine other mammalian species. We performed a population genomics study of 23 additional reindeer representing both domestic and wild populations and several ecotypes from various geographic locations. We assembled 2.66 Gb (N50 scaffold of 5 Mb) of the estimated 2.92 Gb reindeer genome, comprising 27,332 genes. The results from the demographic history analysis suggested marked changes in the effective population size of reindeer during the Pleistocene period. We detected 160 reindeer-specific and expanded genes, of which zinc finger proteins (n=42) and olfactory receptors (n=13) were the most abundant. Comparative genome analyses revealed several genes that may have promoted the adaptation of reindeer, such as those involved in recombination and speciation (*PRDM9*), vitamin D metabolism (*TRPV5, TRPV6*), retinal development (*PRDM1, OPN4B*), circadian rhythm (*GRIA1*), immunity (*CXCR1, CXCR2, CXCR4, IFNW1*), tolerance to cold-triggered pain (*SCN11A*) and antler development (*SILT2*). The majority of these characteristic reindeer genes have been reported for the first time here. Moreover, our population genomics analysis suggested at least two independent reindeer domestication events with genetic lineages originating from different refugial regions after the Last Glacial Maximum. Taken together, our study has provided new insights into the domestication, evolution and adaptation of reindeer and has promoted novel genomic research of reindeer.

## Introduction

Reindeer (*Rangifer tarandus*) have pivotal economic, societal, cultural and ecological values for indigenous people and pastoralists in northern and subarctic regions of Eurasia. Reindeer are a source of meat, hide and occasionally milk and have been used for transportation. Reindeer were crucial for the colonization of the northernmost parts of Eurasia and have a central symbolic role for the indigenous Sami, Nenets, and Evenki cultures and several other North Eurasian cultures^1,2^.

During the evolutionary time scale and in recent demographic history, reindeer were distributed across the northern Eurasian regions, except during the glacial periods^3^, and adapted to a challenging northern environment characterized by short daylight and limited vegetation for feeding during the long winter and prolonged daylight during the short summer period^4,5^. The marked fluctuations in daylight may have led to weakened circadian rhythms^6–8^. Similarly, reindeer exhibit retinal structural adaptation to the extreme seasonal light change to increase retinal sensitivity in dim light^9^. Moreover, reindeer have developed unique features, such as fat metabolism processes, a low resting metabolic rate^5^ and the annual growth and loss of antlers both in males and females^10^.

The genetic diversity, population structure, and prehistorical demographic events of *Rangifer* populations have been investigated, and inferences of domestication have been drawn using mitochondrial DNA (mtDNA) D-loop polymorphisms and autosomal microsatellites as genetic markers^3,11–14^. For example, mtDNA and microsatellite diversity patterns between domestic and wild Eurasian tundra reindeer (*R. t. tarandus*), domestic and wild boreal forest reindeer (*R. t. fennicus*), and Arctic Svalbard reindeer (*R. t. platyrhynchus*), wild North American Caribou (*R. t. caribou*) and several other populations appear to reflect their refugial origins and colonization of circumpolar regions after the Last Glacial Maximum rather than following the classic division of *Rangifer* populations into three main ecological groups adapted to a particular set of environmental conditions (forest, tundra and arctic environments) and sedentary or migratory life-history strategies^3,13,15^. Moreover, mtDNA analyses in ancient and modern *Rangifer* populations indicate at least three different domestication events in the prehistory of reindeer and suggest that one of the domestication sites previously assumed may not have been located in northern Fennoscandia^13,15^.

For most domesticated animal species (e.g., chicken^16^, taurine cattle^17^, pig^18^, sheep^19^ and horse^20^), several genomic tools, such as annotated *de novo* genome assemblies, resequencing data sets, and whole-genome single nucleotide polymorphism (SNP) panels, are available for genomics studies. These whole-genome research approaches have provided critical, new knowledge of the evolution, domestication, genetic diversity, selection patterns and adaptation of domestic animal species. For reindeer genetics studies, however, these genomics tools have become available only recently: Li et al. (2017)^21^ reported the *de novo* sequencing of one female reindeer (*R. tarandus*) originating from Inner Mongolia, PR of China. Subsequently, the reindeer reference genome was compared with the genomes of 50 additional ruminant species, and the genomic data from three Chinese domestic reindeer and three Norwegian wild tundra reindeer were resequenced in the study by Lin et al. (2019)^4^. The genetic basis of specific *R. tarandus* characteristics, such as adaptation to marked annual regulation in daylight, exceptional vitamin D metabolism and behavioural traits, was identified. In the present study, we provide an alternative draft reference genome for reindeer studies: we deep-sequenced and *de novo* assembled the genome of a male reindeer originating from Sodankylä, northern Finland by using the Illumina HiSeq platform. In addition, we analysed the whole transcriptome of six tissues of the reference animal to improve the annotation. Moreover, we resequenced 23 *Rangifer* from Norway and Russia (wild and domestic), together with wild reindeer from Svalbard and Alaska, representing different ecotypes, and report here the largest reindeer genome data sequenced thus far. With these data, we examined the genetic diversity, demographic evolution, adaptation and domestication of reindeer and compared the genome structure of reindeer with that of several other mammalian species, e.g., with several domesticated ruminant species having a *de novo* reference genome sequenced.

## Results

### Genome assembly and annotation

Genomic DNA extracted from the blood sample of a male reindeer was sequenced using the Illumina HiSeq 2500 and 4000 platforms. Libraries with insert sizes ranging from 170 bp to 20 kb were sequenced and generated a total of 513 Gb (170X) raw sequence data (Supplementary Table S1). After filtering low-quality reads, 300.55 Gb (100X) clean data was retained for assembly (Table 1 and Supplementary Table S1). The genome size of reindeer was estimated to be 2.92 Gb using k-mer analysis^22^ (Supplementary Table S2 and Supplementary Fig. S1). High-quality reads were assembled using SOAPdenovo V2.04^23^, and an assembly spanning 2.66 Gb of an estimated 2.92 Gb was generated, with N50 scaffolds and contig sizes of 5 Mb and 48.8 kb, respectively (Table 1, Supplementary Table S3). The final assembly comprised 23,450 scaffolds (170 kb–28.2 Mb), which represented 2.62 Gb and 107, 910 contigs (200 bp–22.5 kb) that in turn represented an additional 44.9 Mb of the assembled genome. The assemblies covered ∼91% of the estimated reindeer genome size. The GC content of the assembled genome was approximately 41.4% (Supplementary Fig. S2 and Supplementary Fig. S3), which is similar to the GC content of the genomes of other related species^24,25^.

**Table 1.** Assembly and annotation statistics.

| Sequencing | Insert size | Clean data (Gb) | Sequence coverage (X) |
| --- | --- | --- | --- |
| Paired-end libraries | 170, 500, 800 bp | 181.39 | 60.46 |
|  | 2, 5, 10, 20 kb | 119.16 | 39.72 |
|  | Total | 300.55 | 100.18 |
| Assembly | N50 (Kb) | Longest (Kb) | Size (Gb) |
| Contig | 48.8 | 441.2 | 2.62 |
| Scaffold | 5,023.6 | 28,190.2 | 2.66 |
| Annotation | Number | Total length (Mb) | Percentage of genome |
| Genes | 27,332 | 787.3 | - |
| Repeats | - | 1,012.6 | 38.02 |

We evaluated the quality of the draft assembly by aligning high-quality reads from the short insert-size libraries against the assembly using Burrows-Wheeler Alignment (BWA)^26^ and found that the mapping rate was 98.3%. In addition, assembly quality was assessed by aligning the *de novo* assembled transcriptome against the assembly using BLAT^27^. A total of 99% of the assembled transcripts were mapped to the corresponding assembly (Supplementary Table S4). Furthermore, to assess the assembly quality, we also used Benchmarking Universal Single-Copy Orthologs (BUSCO)^28^ genes and found that the assembly contained 94.9% (3,895 of 4,104) complete BUSCOs (Supplementary Table S5).

We predicted a total of 27,332 protein-coding genes with an average of 7.4 exons, 1,305 bp coding sequences (CDS) and 28,808 bp transcripts per gene in the reindeer genome by employing a homology-based and an RNA-assisted approach as described by Curwen et al. (2004)^29^ (Table 1, Supplementary Table S7). In total, 26,838 (98.19%) of the total predicted genes were functionally annotated according to public databases (InterPro, Kyoto Encyclopedia of Genes and Genomes (KEGG), Swiss-Prot, TrEMBL), while a total of 494 (1.81%) genes remained unannotated (Supplementary Table S8). To assess the quality of gene prediction, the gene length distribution, the length of the CDS, exons and introns, and the distribution of exon number per gene were compared with those of the five mammalian genomes in a homology-based prediction. No significant differences in exon length and intron length distribution were observed among the six species (Supplementary Fig. S4). Among the compared species, the gene models derived for reindeer were similar to those for dog with respect to all of these key parameters (Supplementary Fig. S4). The annotated gene set was evaluated by BUSCO, and our gene set mapped 96.8% (3,973 of 4,104) complete BUSCOs (Supplementary Table S6), indicating the high quality of our draft genome annotation. We also annotated 6,580 non-coding RNAs (ncRNAs), of which 551 were microRNAs (miRNAs), 329 were rRNAs, 3,994 were tRNAs, and 1,706 were small nuclear RNAs (snRNAs) (Supplementary Table S9). Compared with the analyses of cattle, yak, sheep, goat, horse, human and the first reindeer genome assembly, with less than 900 tRNAs predicted, our analysis revealed an exceptionally high number of tRNAs (the analysis was repeated for confirmation)^21^. This unique finding may be due to some unknown bioinformatics bias or could be related to unique characteristics of the Fennoscandian reindeer.

In total, 38.02% of the assembled reindeer genome comprised repetitive sequences (transposable elements (TEs) and tandem repeats; Table 1, Supplementary Table S10), most of which were transposon-derived repeats. All TEs obtained by different methods (Repbase library^30^, alignment with known transposable-element-related genes, and *de novo* RepeatModeller^31^ identification) were combined, and the result indicated that 36.78% of the reindeer genome consisted of TE sequences (Supplementary Table S11), which is similar to the results obtained for the panda (36.2%)^24^ and dog genomes (36.1%)^32^, while lower than the results obtained for the cattle (46.5%)^17^, goat (42.2%)^33^ and sheep genomes (42.7%)^19^.

### Evolution of the reindeer genome

We compared the novel reindeer reference genome to nine different mammalian species (*Camelus dromedarius, Capra hircus, Ovis aries, Bos taurus, Bos grunniens, Equus caballus, Canis familiaris, Ursus maritimus* and *Homo sapiens*) to examine the evolution of genes and gene families. The *R. tarandus* genome revealed 17,709 orthologous gene families that were shared by at least two species (pairwise comparison to reindeer) and a total of 11,640 orthologous gene families that were shared by all ten species (Fig. 1A, Supplementary Table S12).

**Fig. 1.**
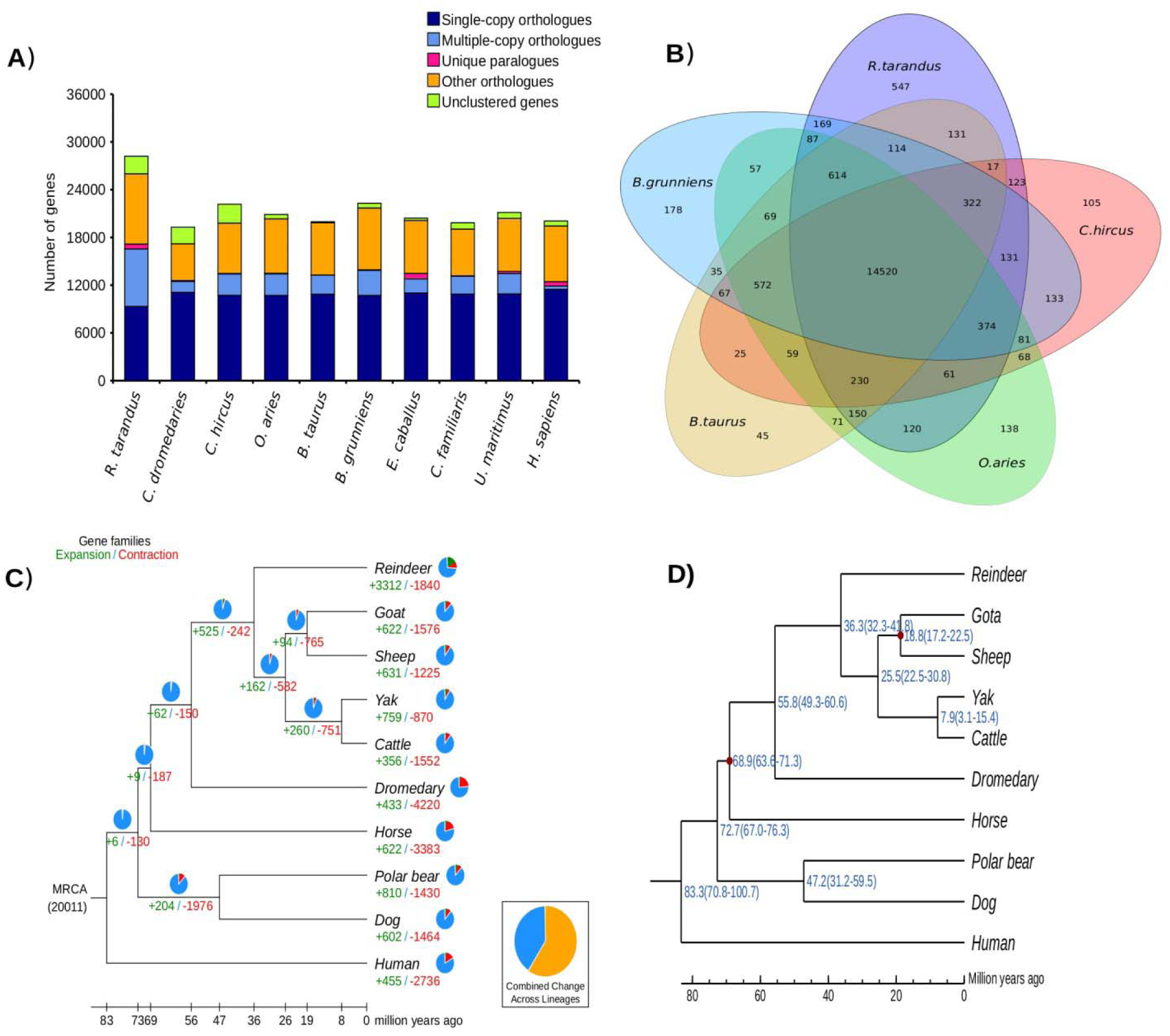
Gene ontology comparison among genomes of reindeer and nine mammalian species. **A)** Distribution of gene clusters among the compared mammalian species. **B**) Venn diagram showing unique and shared gene families among five ruminants (cattle, goat, reindeer, sheep and yak). **C**) Gene expansion and contraction in the reindeer genome. The numbers of gene families that have expanded (green, +) and contracted (red, -) after splitting are shown on the corresponding branch. **D**) Phylogenetic tree based on 4-fold degenerate sites of 7,951 single-copy orthologous genes. Estimates of divergence time with 95% CI are shown at each node.

#### Reindeer-specific gene families

A total of 14,520 orthologous gene families were shared between the domestic ruminant species, while 547 gene families were specific to reindeer (Fig. 1B, Supplementary Data 1). The reindeer-specific gene families contained 1,090 genes, of which 530 were associated with 945 known InterPro domains (Supplementary Data 2). A number of olfactory receptors (n=29 genes), zinc finger proteins (n=71 genes) and ribosomal proteins (n=29 genes) were present in the reindeer-specific gene families (Supplementary Data 1). Out of 1,090 reindeer-specific genes, 691 genes lacked Gene Ontology (GO) annotations. Analysis of the remaining 399 genes revealed 39 significant (false discovery rate (FDR) < 0.05) GO terms, of which 23 were associated with biological processes (BP) and the rest were categorized in molecular function (MF) (Supplementary Table S13). The specific gene families enriched in biological processes revealed several GO terms associated with biosynthetic processes (n=10 GO terms) and metabolic processes (n=7 GO terms) (Supplementary Table S13). The biological processes associated with reindeer-specific gene families also included “GO:0097659, nucleic acid-templated transcription, P=6 × 10^-6^”, “GO:0010468, regulation of gene expression, P=6.2 × 10^-6^” and “GO:0007186, G protein–coupled receptor signaling pathway, P=0.00028” (Supplementary Table S13). Similarly, the molecular function category of the GO terms revealed a number of receptor activities (n=7 GO terms), such as “GO:0099600, transmembrane receptor activity, P= 3.3 × 10^-6^”, “GO:0004888, transmembrane signaling receptor activity, P= 1.8 × 10^-5^”, “GO:0004984, olfactory receptor activity, P= 5.9 × 10^-5^”, “GO:0038023, signaling receptor activity, P= 6.5 × 10^-5^” and “GO:0001637, G protein–coupled chemoattractant receptor activity, P=2.9 × 10^-5^” (Supplementary Table S13). Moreover, three molecular function categories associated with NADH dehydrogenase activity were significantly enriched in reindeer-specific gene families (Supplementary Table S13).

We further examined reindeer-specific gene families to search for genes related to adaptation to subarctic climatic and other environmental factors, such as cold tolerance, marked fluctuations in daylight and feed shortage during winter. Several genes (*AKAP5, CACNB2, CEBPB, CEBPE, COL1A2, CYSLTR1, DNAJB6, EIF2B2, FOXE3, GATA3, GNAO1, MICB, PKD1, PRDM7, PRDM9, RPS6, TRPV5* and *TRPV6*) (Supplementary Data 1) from our list of reindeer-specific gene families have been reported to be associated with cold adaptation in Siberian human populations^34^. We also identified genes related to unique characteristics of the reindeer species^4^, such as vitamin D metabolism (*TRPV5* and *TRPV6*) and antler development (*SILT2*). In addition, we found that the melanopsin gene (*OPN4B*), which is a photoreceptor, has an important function in the retina^35–37^.

#### Expanded and contracted gene families

Based on the comparison of orthologous gene families among the 10 species, the reindeer genome revealed 3,312 and 1,840 expanded and contracted gene families, respectively (Fig. 1C). The significantly (P<0.01) expanded (n=368) and contracted (n=15) gene families contained 2,683 and 19 genes, respectively (Supplementary Data 3 and Data 4). A high number of expanded genes were associated with ribosomal proteins (n=658 genes), zinc finger proteins (n=155 genes) and olfactory receptors (n=35 genes) (Supplementary Data 3). We performed functional enrichment analysis for only the 368 expanded (2,688 genes) gene families (Supplementary Table S14). The expanded genes revealed 163 significant (FDR <0.05) GO terms, of which 122 were categorized into biological processes and 41 were associated with molecular function (Supplementary Table S14 and Data 5). The expanded gene families enriched in biological processes were mainly associated with metabolic processes (n=33 GO terms), biosynthetic processes (n=23 GO terms), ion homeostasis (n=14 GO terms) and transport (n=14 GO terms) (Supplementary Table S14 and Data 5). Similarly, the expanded gene families enriched in the molecular function category were dominated by binding activities (n=16 GO terms), such as “GO:0008199, ferric iron binding, P=4.2 × 10^-57^”, “GO:0005506, iron ion binding, P=2.5 × 10^-17^”, “GO:0031492, nucleosomal DNA binding, P=9.2 × 10^-13^” and “GO:0043566, structure-specific DNA binding, P=9.2 x×10^-13^”.

We further explored expanded gene families to search for genes related to arctic and subarctic adaptation and unique reindeer characteristics. Several genes (*ADRA2A, ADRA2B, ADRA2C, CIDEA, CRYAB, CYCS, DNAJA1, DRD3, EPHA3, EPHB1, FOXC1, FOXC2, GNG5, HMGN3, HOXA5, HSF1, HSPE1, HTR1B, HTR2A, PARK2, PPP2R1A, PRDM1, PRDM7, PRDM9, RAP1B, RHOC, RPS6, SLC25A5, SLC8A1, TRPV5* and *TWIST1)* (Supplementary Data 3) that were predicted to be in expanded gene families have been reported to be associated with cold adaptation in Siberian human populations^34^. Moreover, we also identified genes related to vitamin D metabolism (*TRPV5*) among the list of expanded gene families^4^.

#### Genes under positive selection in reindeer

We conducted an analysis of the signature of positive selection by assessing dN/dS ratios of 7,951 single-copy orthologous gene sets found in the genomes of *R. tarandus* and the other nine mammalian species. A total of 418 of the 7,951 (5.26%) single-copy orthologous genes in reindeer revealed a significant signature of positive selection (P<0.01) (Supplementary Data 6). Analysis of the GO enrichment for the genes under positive selection in reindeer revealed seven statistically significant (FDR ≤ 0.05) enriched GO terms particularly related to channel activities (n=6 GO terms) (Supplementary Table S15).

A number of genes (*BDKRB2, CORIN, GCG, GNB1, IL6, KCNMA1, KCNMB2, MED1, MLXIPL, NPY, NTS, PRDM10, PRKCQ, RNPEP, SHC1, SORBS3, STRADA, TCF7L2, TRPC1* and *VEGFA*) (Supplementary Data 6) showing positive selection in reindeer have been reported to be associated with cold adaptation in Siberian human populations^34^. Moreover, we noticed that a gene regulating circadian rhythm (*GRIA1*)^4^ showed signs of positive selection.

#### Evolutionary divergence analysis

We also estimated the evolutionary divergence of reindeer by comparing the assembled reindeer genome with four other domesticated ruminant species (goat, sheep, yak and taurine cattle), dromedary camel, horse, polar bear, dog and human. A phylogenetic tree was constructed based on 4-fold degenerate codon sites extracted from 7,951 single-copy orthologous genes identified by TreeFam^38^. The estimated divergence time showed that reindeer shared a common ancestor with other domesticated ruminant species approximately 36 million years ago (95% CI 32-42 million years ago) (Fig. 1D). Our divergence estimate is longer than the divergence estimate from a previous reindeer study^21^. Our analysis suggests that the divergence between the domesticated ruminant species and the dromedary camel occurred in the early phase of *Artiodactyla* evolution^39^, 56 million years ago.

### Demographic history

We applied the pairwise sequentially Markovian coalescent (PSMC)^40^ method to the reindeer reference genome to examine the changes in the effective population size (N_e_) of the ancestral population of the Fennoscandian reindeer in the course of the Pleistocene period in the world’s history (Fig. 2). The N_e_ of the ancestor of reindeer gradually decreased between 1 million years ago and 500 thousand years ago (Kya). The ancestral N_e_ of reindeer showed peaks at 150 Kya and 20 Kya, while the population underwent three major bottlenecks at 600 Kya, 40 Kya and 11 Kya.

**Fig. 2.**
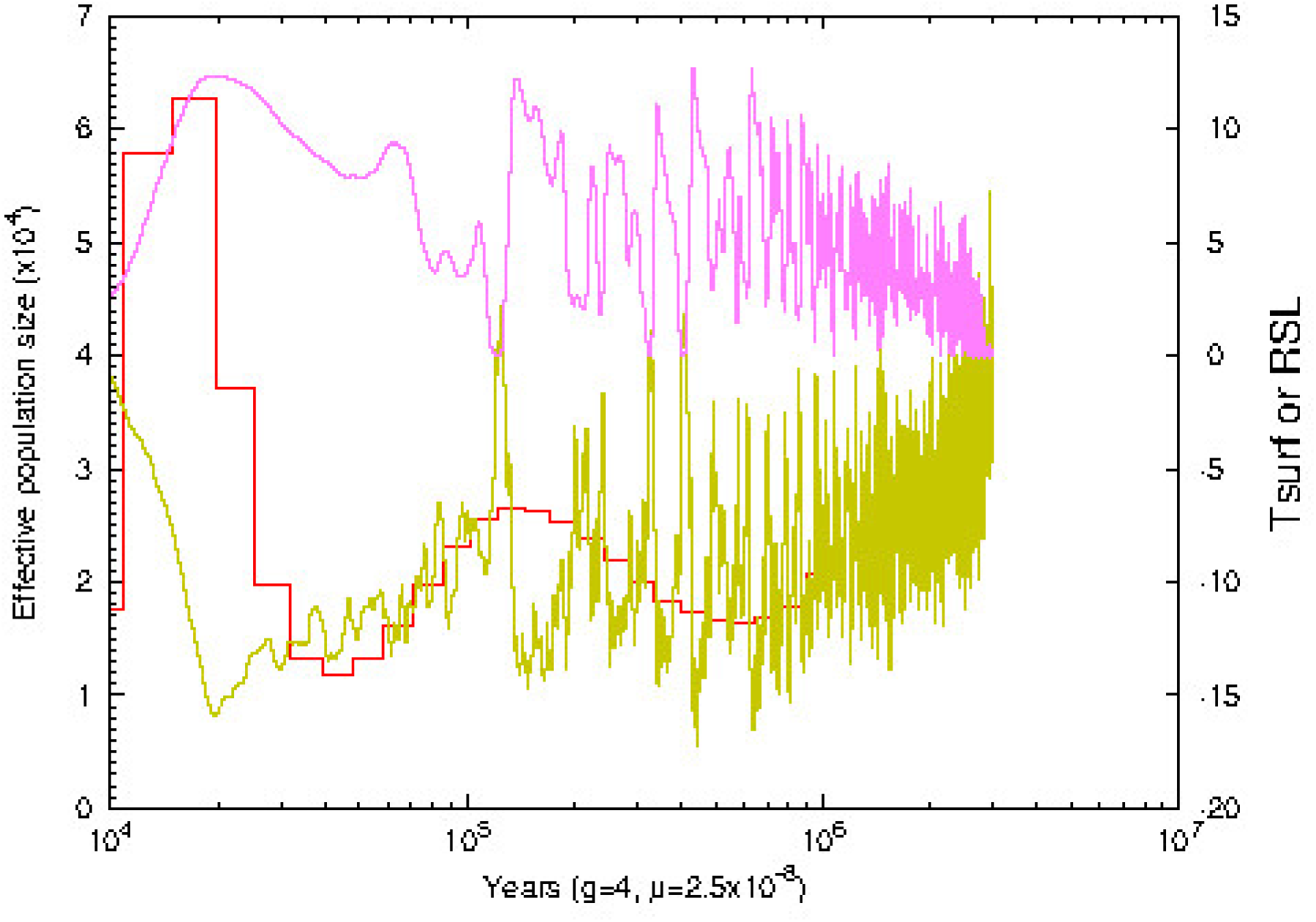
Demographic history of *R. tarandus* reconstructed from the draft reference genome by using PSMC. The X axis shows the time in thousands of years (Kyr), and the Y axis shows the effective population size.

### Heterozygosity estimation

Using the assembled draft reindeer genome sequence as a reference, we mapped the reads of short insert-size libraries to the genome assembly of reindeer and found high coverage, ensuring a high level of accuracy at the nucleotide level. Approximately 98.34% of the reads were successfully mapped to the genome, and the sequencing coverage was approximately 60X. We used the unique map reads to call variants using the Genome Analysis Toolkit (GATK)^41^ best practice pipeline. We identified a total of 5.46 million (M) heterozygous SNPs in the genome. The estimated heterozygosity rate of reindeer was 2.05 × 10e-3, which is 3.48 and 2.3 times higher than that of cattle (0.59 × 10^-3^) and yak (0.89 × 10^-3^)^25^.

### Whole-genome sequence analysis

The raw sequences for the 23 reindeer samples were generated at Beijing Genomics Institute (BGI) using the Illumina HighSeq 4000 platform. The raw reads were preprocessed; adapters and low-quality data were removed. After the data were processed, we generated a total of 680 Gb clean paired-end resequence data (Supplementary Table S16). On average per individual, we achieved 197 Mb and 29.57 Gb clean reads and bases, respectively (Supplementary Table S16.) On average per individual, 98.78% of the clean reads were successfully mapped to the reindeer genome assembly (Supplementary Table S17) and represented 9.46-fold coverage.

A total of 28.6 M high-quality SNPs were detected in the mapped reads across all 23 individuals. The average number of SNPs detected per individual was 7.24 M (Supplementary Table S18). In the wild tundra reindeer in Russia (four samples collected at two sites), 8.00 M SNPs were identified on average per individual; in the wild Norwegian tundra reindeer, 7.09 M (five animals) SNPs were identified; and in the domestic tundra reindeer, 6.79 M (four animals from two sites) SNPs were identified. Three arctic reindeer samples from Svalbard Norway showed low diversity, with an average of 1.37 M (20.42%) heterozygous and 5.35 M (79.58%) homozygous SNPs per individual, while in the rest of the samples, the respective estimates were M (52.14%) and 3.50 M (47.86%). We also detected a total of 3.93 M indels across all 23 samples, and on average, we detected approximately 895,386 indels per individual.

#### Functional annotation of SNPs

SnpEff^42^ was used to annotate all filtered SNPs identified across all samples, and the proportions of SNPs in the intergenic, intronic, downstream and upstream, and transcript regions were 58.27%, 14.0%, 12.52% and 14.61%, respectively, while a small number of variants were annotated in exonic (0.57%) regions (Supplementary Table S19 A and B). The overall estimated missense to silent ratio for all samples was 0.7963. The average SNP-variant rate was 1 variant every 90 bases for all samples. The transition to transversion (Ts/Tv) ratio was calculated for each sample, and the average Ts/Tv ratio was 1.97 for all reindeer samples, which is slightly lower than that of human (2.1)^43^ and lower than that of bovine (2.2)^44–46^.

#### Population genetic analysis

Genetic relationships between the 23 resequenced animals and the reference animal were investigated with a principal component analysis (PCA). The PCA plot revealed two major groups Fennoscandia reindeer versus forest and tundra reindeer/caribou from Russia and Alaska, as shown in Fig. 3. The reindeer from Svalbard and Novaja Zemlya, both wild arctic ecotype, grouped separately from the main clusters. We computed genetic diversity parameters for the two main clusters. The overall genome-wide genetic diversity, as measured by Watterson’s (θ) and pairwise nucleotide diversity (π), was smaller in the Fennoscandian group (0.002022733 and 0.002096655, respectively) than in the Russian – Alaskan group (0.002441424 and 0.002265819, respectively). The genetic differentiation between the two main groups, as measured by F_st_, was 0.039786, suggesting that these two cluster populations exhibit low differentiation.

**Fig. 3.**
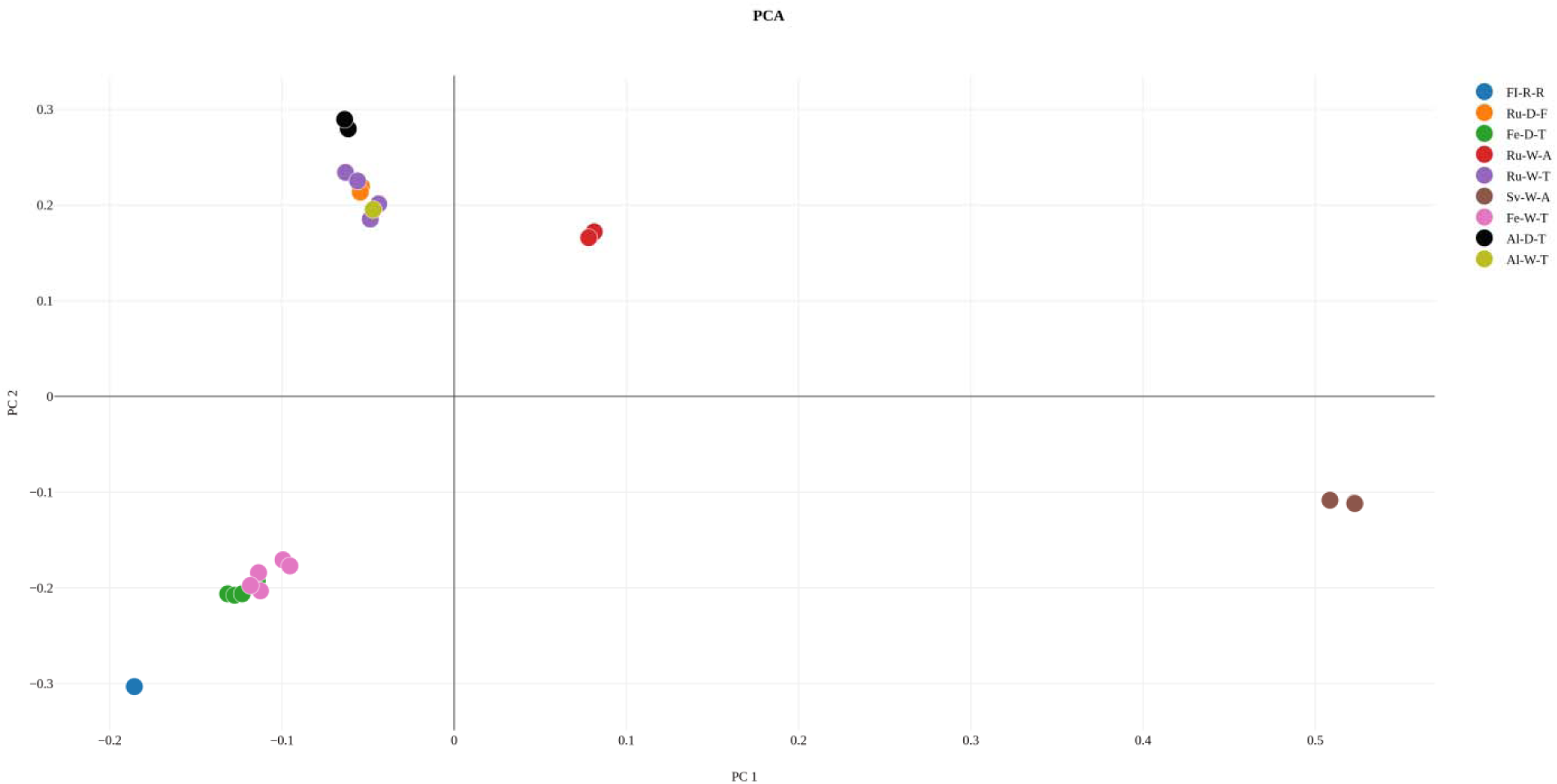
PCA of 23 resequenced animals and the reference individual. PCA plot. (FI-R-R, Finnish reindeer (i.e. the reference animal); Fe-W-T, Fennoscandia wild tundra reindeer; Fe-D-T, Fennoscandia domestic tundra reindeer; Ru-W-T, Russian wild tundra reindeer; Ru-D-T, Russian domestic tundra reindeer; Ru-D-F, Russian domestic forest reindeer; Ru-W-A, Russian wild arctic reindeer (i.e. Novaia Zemlia); Sv-W-A, Svalbard wild arctic reindeer; Al-D-T, Alaska Domestic tundra reindeer; Al-W-T, Alaska wild tundra caribou)

### Assembly and annotation of the reindeer mitochondrial genome

The reindeer mitochondrial genome was assembled using the mitochondrial baiting and iterative mapping (MITObim)^47^ approach. A total of 16,962 reads were utilized in the assembly, corresponding to 0.036% of the reads generated from the 170 bp insert-size library. The reads used in the assembly mapped with 84% of bases mapping with quality >Q20. The assembled mitochondrial genome is 16,357 bp in length with an average conversion of 131.84X and a GC content of 36.28%. The reindeer mitogenome consists of 22 tRNAs, 13 protein-coding genes and two rRNAs (Supplementary Table S20).

## Discussion

With our present study and the recent publications by Li et al. (2017)^21^ and Lin et al. (2019)^4^, a new genomic era in the study of reindeer has been established. Here, we have generated and described an alternative draft genome assembly for reindeer and published the largest resequenced *Rangifer* sp. dataset for the investigation of genetic diversity, domestication, demographic aspects and genomic adaptation.

Our reindeer genome assembly is 2.66 Gb in size and represents ∼91% of the estimated 2.92 Gb genome size according to the k-mer analysis. The estimated genome size of reindeer in our study is slightly higher than that in two recent studies, i.e., 2.86^48^ Gb and 2.76 Gb^21^. Species belonging to the *Cervidae* family^48^, such as Indian muntjac (2.92 Gb), Chinese muntjac (2.99 Gb), Milu (3.09 Gb), Black muntjac (3.24 Gb) and White-lipped deer (3.23 Gb), appear to have larger genome sizes than domesticated mammalian species in general (∼2.6–2.7 Gb), such as pig (*Sus scrofa*^18^), yak (*B. grunniens*^25^), dromedary camel (*C. dromedarius*^49^) and sheep (*O. aries*^19^). In addition, the number of protein-coding annotated genes (>27,000) in our genome assembly draft is higher than that typically predicted in the latest assemblies of several other domesticated mammalian species (∼21,000)^18–20,25,33^ and in the first genome draft of reindeer (21,555)^21^. We assume that the differences in the genome size and the gene numbers may be due to the annotation approach (homology-based approach) used in this study and will require further validation. However, the BUSCO assessment revealed 97% orthologous genes in the present assembly, which is better than the BUSCO assessment results in a previous study^21^, thus indicating the high quality of our reindeer gene annotation. In addition, the key gene parameters (see Supplementary Table S7 and Fig. S4) also revealed no significant differences between the reindeer genome and the five species that were used in homology predication related to gene and exon length distribution.

We applied the present draft assembly to estimate the evolutionary divergence times between reindeer and nine other mammalian species and to reveal recent demographic events in the history of reindeer that occurred during the last 1 million years. Our phylogeny suggests that reindeer diverged from four other domestic ruminants ca. 36 million years ago, which is earlier than previously estimated ^21,48^. The demographic analysis (Fig. 2) showed that there have been marked fluctuations in the N_e_ of the reindeer species: two population expansions and three bottlenecks. The decline in the N_e_ occurred during the mid-Pleistocene period, and the peak during Last Glacial Maximum period may have been associated with global temporal changes in climate ^50–52^. The pattern of the N_e_ of the reindeer genome during the Pleistocene period is consistent with the previously suggested hypotheses of the responses of other domestic mammalian species, such as pig (*S. scrofa*^18^), horse (*E. caballus*^53^), sheep (*O. aries*^54^) and cattle (*B. taurus*^55^), to temporal climate changes in the past. However, the heterozygote rate (2.05 × 10^-^ 3) of the reindeer genome estimated in our analysis was 3.48 and 2.3 times higher than that of *B. taurus*-cattle and *B. grunniens*-yak, respectively^25^, suggesting a larger founder population size of the contemporary semi-domesticated reindeer. In addition, possible gene flow from wild reindeer populations, a less intensive artificial selection history and the early phase of reindeer domestication history in general may have contributed to higher diversity estimates for reindeer than for the two other ruminant species^15^.

To explore the genetic distinctiveness of reindeer in the present research context, we focused on reindeer-specific and expanded orthologous gene families and their genes and gene functions. The changes in genetic makeup (i.e., gene copy and/or protein domain number changes) are potentially linked to unique phenotypic and functional changes, suggesting a mechanism for adaptive divergence of closely related species^24,25,56^. Gene family analysis revealed that 160 genes were shared between 547 reindeer-specific gene families and 368 expanded gene families. Therefore, the 160 genes that we identified as unique to reindeer compared to four other ruminants are also rapidly evolving (among nine other mammals) and may thus represent characteristic features of reindeer. Zinc finger proteins (n=42) dominated the list of uniquely expanding genes followed by olfactory receptors (n=13). Zinc finger proteins have multi-functional roles, including lipid binding and the differentiation of adipose tissues^57,58^. Interestingly, among the list of zinc finger proteins was PR domain zinc finger protein 9 (*PRDM9*), which is considered one of the fastest evolving genes in mammalian species. *PRDM9*, also referred to as the speciation gene, has an important function in enhancing recombination and epigenetic modification^59,60^. We speculate that the *PRDM9* gene has played a vital role in the evolution and adaptation of *R. tarandus* to challenging northernmost biogeographic and climatic conditions. Another member of the zinc finger protein, PR domain zinc finger protein 1 (*PRDM1*), is associated with retinal development^61–63^ and is a candidate gene promoting adaptation to extreme seasonal light changes. Furthermore, the improved sensory systems of reindeer were also reflected in our study, where we observed several genes associated with olfactory receptors and G protein–coupled receptors in specific and expanded gene families (Supplementary Data 1 and Data 3). The mammalian genome contains 4–5% of olfactory receptor genes, which are sensors to the extracellular environment and essential for the survival of animals^64^. G protein–coupled receptors are also known to be involved in sensing of the extracellular environment^64^. During the long winter season, the reindeer sense of smell or olfaction has an important role in finding feed covered with snow.

Moreover, the examination of genes in the reindeer-specific and expanded orthologous gene families provided information on genes associated with specific phenotypic characteristics of reindeer. We identified two genes belonging to the transient receptor potential (TRP) cation channel subfamily, *TRPV5* and *TRPV6*, among the reindeer-specific gene families. *TRPV5* and *TRPV6* are involved in calcium reabsorption and maintenance of blood calcium levels in higher organisms^65,66^. Among these, *TRPV5* has been found to be associated with vitamin D metabolism^4^. Previous studies have reported that reindeer require efficient calcium metabolism and reabsorption for antler growth during the period of continuous winter darkness and low solar energy^67,68^. Studies also indicated that efficient vitamin D metabolism is needed to sustain calcium metabolism and reabsorption^67,68^. The nucleotide sequences of both the *TRPV5* and *TRPV6* genes harbour 15 exons, encode proteins of approximately 730 amino acids and have 75% sequence similarity^65,66^. Both genes have similar functional properties and regulation mechanisms^65^. We speculate that in addition to *TRPV5, TRPV6* also has a significant role in vitamin D metabolism in reindeer. In addition, three adaptive immune-related chemokines (*CXCR1, CXCR2, CXCR4*)^69–71^ and three interferons (*IFNT1, IFNT3, IFNW1*) that have antiviral functions^72,73^ may have promoted the adaptation of reindeer to the given environment. Finally, insulin-like growth factor 1 receptor (*IGF1R*) promotes the activity of *IGF1*, which has a pivotal role in antler formation (*IGF1* regulates *RUNX1* expression via IRS1/2 signalling for antler growth)^74,75^.

We also observed large numbers of ribosomal proteins present in the reindeer-specific and expanded gene families. A number of reports have suggested that ribosomal proteins are essential for survival and adaptation to environmental stress^76–78^. For example, one of the ribosomal proteins we found was the ribosomal large subunit protein 7 (*RPL7*), which has been previously reported in Russian cattle breeds^79^ to be related to adaptation to harsh (cold) environments. The ribosomal large subunit protein 7 (*RPL7*) gene was found to be upregulated in the skin of freeze-tolerant wood frogs compared to non-tolerant species, showing that this gene is cold-responsive and that the gene was upregulated in skeletal muscle and the brain under freezing stress^78^. Moreover, in the list of reindeer-specific gene families, we found opsin 4B (*OPN4B)*, which is a photoreceptor that appears to be involved in the regulation of the circadian clock^35–37^. Several studies^37,80–82^ have reported the existence of a melanopsin-associated photoreceptive system in the mammalian retina that helps to transmit photic information and is also involved in the regulation of photoentrainment of the circadian clock. A previous study^9^ reported that reindeer possess a tapetum lucidum (reflective surface behind the central retina) to cope with winter darkness, which may help in changing wavelength reflection and increase the sensitivity of the animal’s vision in dim light. In winter, the reindeer tapetum lucidum changes to blue, which may help scatter light through photoreceptors, and less light is reflected^9^. We speculate that this melanopsin-related gene might have important functions for the adaptation to the light system in circumpolar environments. Furthermore, we found Slit homolog 2 protein (*SLIT2*) in specific gene families, which has been reported to be associated with antler development in a recent reindeer study^4^.

In our study, signs of positive selection were found in genes in the genome of *R. tarandus* mainly enriched in GO terms related to channel activity (Supplementary Table S15), such as sodium channel activity and ion channel activity, which are thought to be relevant in cold sensing mechanisms^83^. The ability to detect and adapt to cold temperature is crucial for the survival of an organism^84,85^. The process of sensory transduction and a large array of ion channels, such as Na^+^ and K^+^ channels, are involved in cold temperature detection^83^. Among the genes categorized into these GO terms, interestingly, *SCN11A1* has an essential role in pain tolerance caused by cold stress^86,87^, suggesting the gene’s role in survival in extremely cold environments. One additional gene showing signatures of positive selection, glutamate ionotropic receptor AMPA type subunit 1 (GRIA1), appears to have a role in the circadian clock^4^.

Finally, we investigated the genetic relationships between the *de novo* reference animal and the 23 resequenced individuals, and two main genetic clusters (Fennoscandia and Russia-Alaska) were identified in our data (Fig. 3). We suggest that this observation reflects the historical spread of reindeer populations in northern Eurasia and reveals the domestication history of reindeer. The Russian/northern American cluster probably reflects the Euro-Beringian lineage evolved from Pleistocene population in northern Eurasia^3,88^ while the Fennoscandia cluster may have descended from some refugia populations in southern Europe partly isolated from the Euro-Beringian lineage^89^. In these two main genetic clusters, we found both domestic and wild reindeer from respective Fennoscandia and Russia, which gives support to previously indicated independent domestication origin for the breeds in these regions^13,90^. Our findings are consistent with the previous study^15^. Our data suggest that the semi-domesticated tundra reindeer show lower genetic diversity in terms of the number of SNPs per individual than the wild tundra reindeer, implying a genetic bottleneck effect caused by domestication as also seen in temporal change in mitochondrial DNA^13,91^. Moreover, our data clearly indicated the effects of geographic isolation and genetic bottleneck in the Svalbard reindeer: the subspecies (or an arctic ecotype) displayed exceptionally low diversity compared to the other resequenced Rangifer populations.

Here, we have reported a 2.66 Gb genome assembly of a Finnish male reindeer originating from Sodankylä (67.34°N, 26.83°E), where the annual mean temperature is 0 °C (the mean temperature in January and February -13 °C), the snow cover is on average 202 days per year, the daylight is less than 3 h from December until mid-January (during a short period in December, the sun is down all day) and the sun is up all day from June until mid-July (https://www.timeanddate.com/). The first draft assembly of the reindeer genome was obtained from a female animal^21^ from Inner Mongolia Autonomous Region, China (50.77°N, 121.47°E), where the climate is continental, but the annual daylight rhythm (at least 8 h of daylight and no periods of total darkness or “nightless nights” exist) is different from that typically found in traditional reindeer herding regions in northern Eurasia. In addition to these climatic differences in their geographic origins, we assume that the Finnish reindeer and Inner Mongolian reindeer do not share their genetic origins in the same refugial, ancestral populations. The Finnish reference reindeer was found to belong to the cluster of northern Fennoscandia reindeer distinct from the “eastern” and North American genetic cluster. It is recommended that phylogenetically different animal populations have their own reference assemblies^55^. With our GO family-based comparisons with nine other mammalian genomes, we identified hundreds of genes and gene families that are unique, under positive selection and rapidly expanding in reindeer. Exploring these genes and gene families revealed several reindeer-specific characteristics that have helped these animals survive in the arctic and subarctic conditions. We identified genes that are important for vitamin D metabolism (*TRPV5, TRPV6*), antler formation (*SLIT2*), and circadian rhythm (*OPN4B*). While many of the findings from our study complement reports from other species, the functions of *TRPV6* and *OPN4B* have been reported here for the first time. Population genomics analyses based on 23 individuals representing domestic and wild reindeer suggested two domestication events in *R. tarandus* subspecies, but additional larger scale studies are required for validating this finding. Collectively, our study provides new insight into the genomic and evolutionary characteristics of reindeer, and the resequencing data (the most comprehensive in reindeer thus far) will serve as an invaluable resource for future studies involving reindeer and other cervids.

## Materials and methods

### Sampling and genome sequencing

All protocols and sample collections were performed in accordance with the legislations approved by the Animal Experiment Board in Finland (ESAVI/7034/04.10.07.2015).

A one-year-old male reindeer *(R. tarandus)* from Sodankylä, Finland, was selected, and blood samples were collected for sequencing. Genomic DNA was extracted from the blood using a standard phenol/chloroform method and sequenced at high coverage on the Illumina HiSeq 2500 and 4000 platforms using a shotgun-sequencing approach. We constructed seven paired-end DNA libraries with insert sizes ranging from 170 bp to 20 kb. Summary statistics of the generated reads are shown in Table 1 and Supplementary Table S1. The short reads with low-quality bases were removed before assembly by filtering as follows:

1. Reads from short insert-size libraries having ‘N’ over 2% of its length, and the reads from large insert-size libraries having ‘N’ over 5% of its length.
2. Reads from short insert-size libraries having more than 40% bases with Q20 (quality) less than or equal to 7, and the reads from large insert-size libraries having more than 30% bases with Q20 less than or equal to 7.
3. Reads with adapter sequences.
4. Paired-end reads of read 1 and read 2 that were completely identical.

### Estimation of genome size using k-mer analysis

For the short reads with an insert size of 500 bp, a total of 64.7 Gb (approximately 21.6X) was used to estimate the genome size using the k-mer method^22^, as described in previous studies^24^. We used the following formula to estimate the genome size (G): G=K_num/K_depth, where K_num is the total number of k-mers, and K_depth is the frequency occurring more frequently than other frequencies. In this study, the size of k-mer for reindeer was set to 17, K_num was 55,451,706,382 and K_depth was 19. Hence, the genome size was estimated to be approximately 2.92 Gb (Supplementary Table S2).

### Genome assembly

After the data were filtered, clean data were retained for assembly (Table 1, Supplementary Table S1). The high-quality clean reads were assembled first into contigs and subsequently into scaffolds using SOAPdenovo V2.04 software^23^.

The quality of the genome assembly was evaluated by aligning high-quality reads from short insert-size libraries to the *de novo* genome assembly using BWA^26^ (version. 0.7.15) with no more than five mismatches and then by determining the percentage of total aligned reads. Reads from the transcriptome were assembled by trinityrnaseq-2.0.6^92^; all sequences were mapped to the assembly by BLAT^27^ with the default parameters. We also employed BUSCO (version 3.0.2)^28^ to assess the completeness of the genome assembly and annotated gene set using the conserved 4,104 mammalian genes.

### RNA sequencing

To improve the reference genome annotation, we extracted RNA using the RNeasy Plus Universal Kit (Qiagen, Valencia, CA, USA) from six tissues (scapular adipose, metacarpal adipose, tailhead adipose, perirenal adipose, liver and musculus gluteobiceps) from the same male reindeer subjected to *de novo* sequencing. The tissue samples were collected at slaughter and stored in RNA*later*® Solution (Ambion/Qiagen, Valencia, CA, USA) according to the manufacturer’s instructions. mRNA libraries were sequenced using Illumina HiSeq 2500 with a paired-end (2×150 bp) strategy. The raw data were preprocessed, and adapters and low-quality reads were filtered out using the cutadapt tool^93^.

### Genome annotation

#### Repeat annotation

Tandem repeats were detected in the generated genome assembly using Tandem Repeats Finder (TRF) software^94^. TEs were predicted in the genome using the combination of both known and *de novo* strategies. For the known-based strategy, we identified known TEs in the genome using RepeatMasker^95^ and RepeatProteinMask^96^ by homology search against the Repbase^30^ library with default parameters. For the *de novo* way, we constructed a *de novo* repeat library using RepeatModeler^31^, with the default parameters. RepeatMasker was used again with the *de novo* libraries to identify new TEs in the genome. All repeats obtained by the different methods were combined according to the coordinate in the genome and overlapping TEs belonging to the same repeat class to form a non-redundant list of reindeer repeats.

#### Protein-coding gene prediction

We employed both a homology-based and an RNA-assisted approach to predict protein-coding genes as described by Curwen et al. (2004)^29^. First, we downloaded protein sets from five sequenced animal genomes (*H. sapiens, M. musculus, B. taurus, C. dromedarius and C. familiaris*) from Ensembl (release 89)^97^ and mapped them onto the reindeer genome using tblastn^98^. Second, high-score segment pairs (HSPs) were concatenated between the same pair of proteins by Solar (in-house software, version 0.9.6). Third, homologous genome sequences were aligned against the matching proteins using GeneWise^99^ to define accurate gene models. Then, we filtered redundancy based on the score from GeneWise. Finally, we combined the annotated results from the homologue-based method to form the final genome annotation as described by Curwen et al. (2004)^29^. Functions of genes were assigned based on the best match derived from the alignments to proteins annotated in four protein databases: InterPro^100^, KEGG^101^, Swiss-Prot^102^ and TrEMBL^103^.

#### ncRNA annotation

We used INFERNAL^104^ and tRNAscan-SE^105^ to predict ncRNA. Four types of ncRNAs were annotated in our analysis: tRNA, rRNA, miRNA and snRNA. tRNA genes were predicted by tRNAscan-SE with eukaryotic parameters. rRNA fragments were identified by aligning the rRNA template sequences from human genomes using BLASTN with an E-value cutoff of 1e^-5^. miRNA and snRNA genes were inferred by the INFERNAL software against the Rfam database (v11.0)^106^.

### Gene family construction

To define gene family evolution in the reindeer genome, we used the TreeFam methodology^38^ as follows: BLAST was used to compare all protein sequences from 10 species, *R. tarandus, C. dromedarius, C. hircus, O. aries, B. taurus, B. grunniens, E. caballus, C. familiaris, U. maritimus* and *H. sapiens*, with the E-value threshold set to 1e-7. After that, HSPs of each protein pair were concatenated by Solar software. H-scores were computed based on Bit-scores, and these were taken to evaluate the similarity among genes. Finally, gene families were obtained by clustering homologous gene sequences using Hcluster_sg (version 0.5.0). If these genes had functional motifs, they were annotated by GO^107^.

### Phylogenetic tree construction and divergence time estimation

The phylogenetic tree was constructed based on the single-copy orthologous genes from the ten species identified by gene family analysis. CDS from each single-copy gene cluster were aligned using MUSCLE^108^, and their protein sequences were concatenated to form one super gene for every species. Four-fold degenerate sites (4d sites) of aligned CDS were extracted for subsequent analysis. The phylogenetic relationships were inferred using PhyML software^109,110^, with the GTR substitution model and gamma distribution rates model as the chosen parameters. We constructed a phylogenetic framework including six *Artiodactyla* and four other vertebrates, with *H. sapiens* as the outgroup. Divergence times were estimated using PAML mcmctree (PAML version 4.5)^111–113^ by implementing the approximate likelihood calculation method.

### Expansion and contraction of gene families

We analysed the expansion and contraction of gene families using the CAFE program^114^, which employs a random birth and death model across a user-specified phylogeny. The global parameter λ, which describes both the gene birth (λ) and death (μ = -λ) rate across all branches in the tree for all gene families, was estimated through maximum likelihood. A conditional P-value was calculated for each gene family, and families with conditional P-values less than the threshold (0.01) were considered to have notable gains or losses. Finally, the branches responsible for low overall P-values were defined as significant families.

### Detection of positively selected genes (PSGs)

To estimate positive selection, dN/dS ratios were calculated for all single-copy orthologues of *R. tarandus* and nine other vertebrates (*C. dromedarius, C. hircus, O. aries, B. taurus, B. grunniens, E. caballus, C. familiaris, U. maritimus* and *H. sapiens*). Orthologous genes were first aligned by PRANK^115^, then Gblocks 0.91b was used to remove ambiguously aligned blocks within PRANK alignments and ‘codeml’ in the PAML package^113^ was employed with the free-ratio model to estimate Ka, Ks, and Ka/Ks ratios of different branches. The difference in mean Ka/Ks ratios for single-copy genes between *R. tarandus* and each of the other species were compared with paired Wilcoxon rank-sum tests. The genes that showed values of Ka/Ks higher than 1 along the branch leading to *R. tarandus* were reanalysed using the codon-based branch-site tests implemented in PAML. The branch-site model, which allowed ω to vary both at sites in the protein and across branches, was used to detect episodic positive selection.

### SNP calling and heterozygosity estimation

We used high-quality reads from the short insert-size libraries (170, 500 and 800 bp) to call SNPs. The quality reads were mapped against the assembly using BWA^26^ (version. 0.7.15) software with the default parameter. After mapping, we employed Picard tools (http://broadinstitute.github.io/picard/) (version.2.5.0) to preprocess alignment reads and remove PCR duplicates, and then the uniquely mapped reads were used for SNP calling. Further, we used GATK (version. 3.6)^41^ to identify poorly mapped regions nearby indels, realigned and performed SNP calling according to the GATK best practice pipeline^116^. We estimated the heterozygosity rate of the reindeer genome; first, we counted the identified heterozygous SNPs and then divided the total heterozygous SNPs by the effective genome size.

### Population history estimation

We inferred a marked population bottleneck in the demographic history of *R. tarandus* using the pairwise sequentially Markovian coalescent (PSMC) model^40^ with generation times and mutation rates.

### Assembly and annotation of the reindeer mitochondrial genome

Complete mitochondrial genomes are useful materials for phylogenetic studies. The complete mitochondrial genome for reindeer was assembled using MITObim v 1.9 software^47^. Among the reads generated from the seven libraries on the reference individual, for the mitochondrial genome assembly, we used only reads from the 170 bp insert-size library (10% of 66.2 Gb, Table S1) for MITObim input. The short DNA sequence (506 bp), cytochrome oxidase subunit 1 (COI) (GenBank accession number COI: KX085230.1)^12^ from *R. tarandus* was used to seed the initial baiting of mitochondrial reads. Following the assembly, the *de novo* assembly was annotated using the MITOs web server^117^ with the default parameters.

### Whole-genome resequencing and variant calling of 23 domestic and wild reindeer

To investigate genome variation and perform population analyses, we resequenced 23 domesticated and semi-domesticated reindeer individuals from Russia, Norway, Alaska and USA. Resequenced samples originated from the following populations and subspecies: domesticated forest reindeer from Russia (n=2), domesticated tundra reindeer from Norway (n=4), wild island tundra-mountain reindeer from Russia (n=2), wild tundra reindeer from Russia (n=4), wild tundra reindeer from Norway (n=5), wild arctic reindeer from Svalbard Norway (n=3), Alaskan domestic reindeer from Alaska-USA (n=2) and Alaskan wild caribou from Alaska-USA (n=1). Genomic DNA was extracted from 23 blood samples (obtained from the previous study by Flagstad and Røed 2003^3^), and sequence libraries with an average fragment size of 500 bp were constructed for each individual. Paired-end reads were generated using Illumina HiSeq 4000 at BGI. After sequencing, the raw reads were filtered to remove adapter sequences, contamination and low-quality reads. Following the data treatment, we achieved an average of 197 Mb and 29.57 Gb clean reads and bases, respectively (Supplementary Table S16.)

The clean reads were mapped against the present draft reindeer assembly genome using BWA with the default parameters. Using Picard tools, we preprocessed and filtered alignments for SNP calling, including the removal of low-quality alignments, the sorting of alignments and the removal of PCR duplicates. Further, using GATK, we identified poorly mapped regions nearby indels from the alignments and performed a realignment and base quality recalibration on the bases that were disrupted by indel sites. Finally, using the GATK best practice pipeline, we performed SNP discovery and genotyping across the 23 samples.

### Functional annotation of SNPs

SnpEff V4.3T^42^ was used to annotate the SNPs and categorize them into coding (synonymous and non-synonymous), upstream/downstream and intronic/intergenic classes. For this analysis, we constructed the SnpEff databases using the draft reindeer reference genome and the corresponding GTF genome annotation file.

### Population genetics statistics

First, to examine the genetic relationships across the 23 reindeer samples, we conducted PCA with smartpca in EIGENSOFT3.0 software^118^ using the detected SNPs in each individual. Second, using the PCA plot result, we divided the 23 samples according to their grouping in the PCA plot and computed the average pairwise nucleotide diversity within a population π and the proportion of polymorphic sites Watterson’s θ for the main clusters using the Bio::PopGen::Statistics package in BioPerl (v1.6.924)^119^. The same program was used to compute the population differentiation estimate, with pairwise F_st_ between the two population groups (i.e., the PCA cluster result).

## Abbreviations

Gb: Gigabases
kb: Kilobases
Mb: Megabases
ncRNA: Non-coding RNA
tRNA: Transfer RNA
rRNA: Ribosomal RNA
miRNA: MicroRNA
snRNA: Small nuclear RNA
BP: Biological process
MF: Molecular function
CDS: Coding sequences
PSMC: Pairwise Sequentially Markovian Coalescent
SNPs: Single nucleotide polymorphisms
BWA: Burrows-Wheeler alignment
GATK: Genome Analysis Toolkit
PCA: Principal component analysis
mtDNA: Mitochondrial DNA
N_e_: Effective population size
Kya: Thousand years ago
M: Million

## Acknowledgements

This study was financially supported by the Academy of Finland in the Arctic Research Programme ARKTIKO (decision number 286040). M. B acknowledges partial funding from the Finnish cultural foundation. We are grateful to Juhani Maijala for providing the reindeer that was used as a reference genome. The authors thank Nuccio Mazzullo, Päivi Soppela and Anna Stammler-Gossmann from the Arctic Centre, University of Lapland, Rovaniemi, Finland for collaboration in the sampling of the Finnish reference reindeer. The authors wish to acknowledge the CSC-IT Center for Science, Finland, for computational resources. The owners of the animals included in the study are acknowledged for providing samples for this study.

## Contributions

J. K conceived and designed the project. J. K, T. R, M. H, J. P and K. R collected the samples. M. W, K. P and M. Y performed bioinformatics analyses. M. W prepared the manuscript draft with substantial contribution from J. K and K.P. All authors read, reviewed and approved the final manuscript.

